# IGF2BP1 is a targetable SRC/MAPK-dependent driver of invasive growth in ovarian cancer

**DOI:** 10.1101/2020.06.19.159905

**Authors:** Nadine Bley, Annekatrin Schott, Simon Müller, Danny Misiak, Marcell Lederer, Tommy Fuchs, Chris Aßmann, Markus Glaß, Christian Ihling, Andrea Sinz, Nikolaos Pazaitis, Claudia Wickenhauser, Martina Vetter, Olga Ungurs, Hans-Georg Strauss, Christoph Thomssen, Stefan Hüttelmaier

## Abstract

Epithelial-to-mesenchymal transition (EMT) is a hallmark of aggressive, mesenchymal-like high-grade serous ovarian carcinoma (HG-SOC). The SRC kinase is a key driver of cancer-associated EMT promoting adherens junction (AJ) disassembly by phosphorylation-driven internalization and degradation of AJ proteins. Here we show, that the IGF2 mRNA binding protein 1 (IGF2BP1) is up-regulated in mesenchymal-like HG-SOC and promotes SRC activation by a previously unknown protein-ligand-induced, but RNA-independent mechanism. IGF2BP1-driven invasive growth of ovarian cancer cells essentially relies on the SRC-dependent disassembly of AJs. Concomitantly, IGF2BP1 enhances ERK2 expression in a RNA-binding dependent manner. Together this reveals a post-transcriptional mechanism of interconnected stimulation of SRC/ERK signaling in ovarian cancer cells. The IGF2BP1-SRC/ERK2 axis is targetable by the SRC-inhibitor saracatinib and MEK-inhibitor selumetinib. However, due to IGF2BP1-directed stimulation only combinatorial treatment effectively overcomes the IGF2BP1-promoted invasive growth in 3D culture conditions as well as intraperitoneal mouse models. In conclusion, we reveal an unexpected role of IGF2BP1 in enhancing SRC/MAPK-driven invasive growth of ovarian cancer cells. This provides a rational for the therapeutic benefit of combinatorial SRC/MEK inhibition in mesenchymal-like HG-SOC.

**Graphical Abstract:** 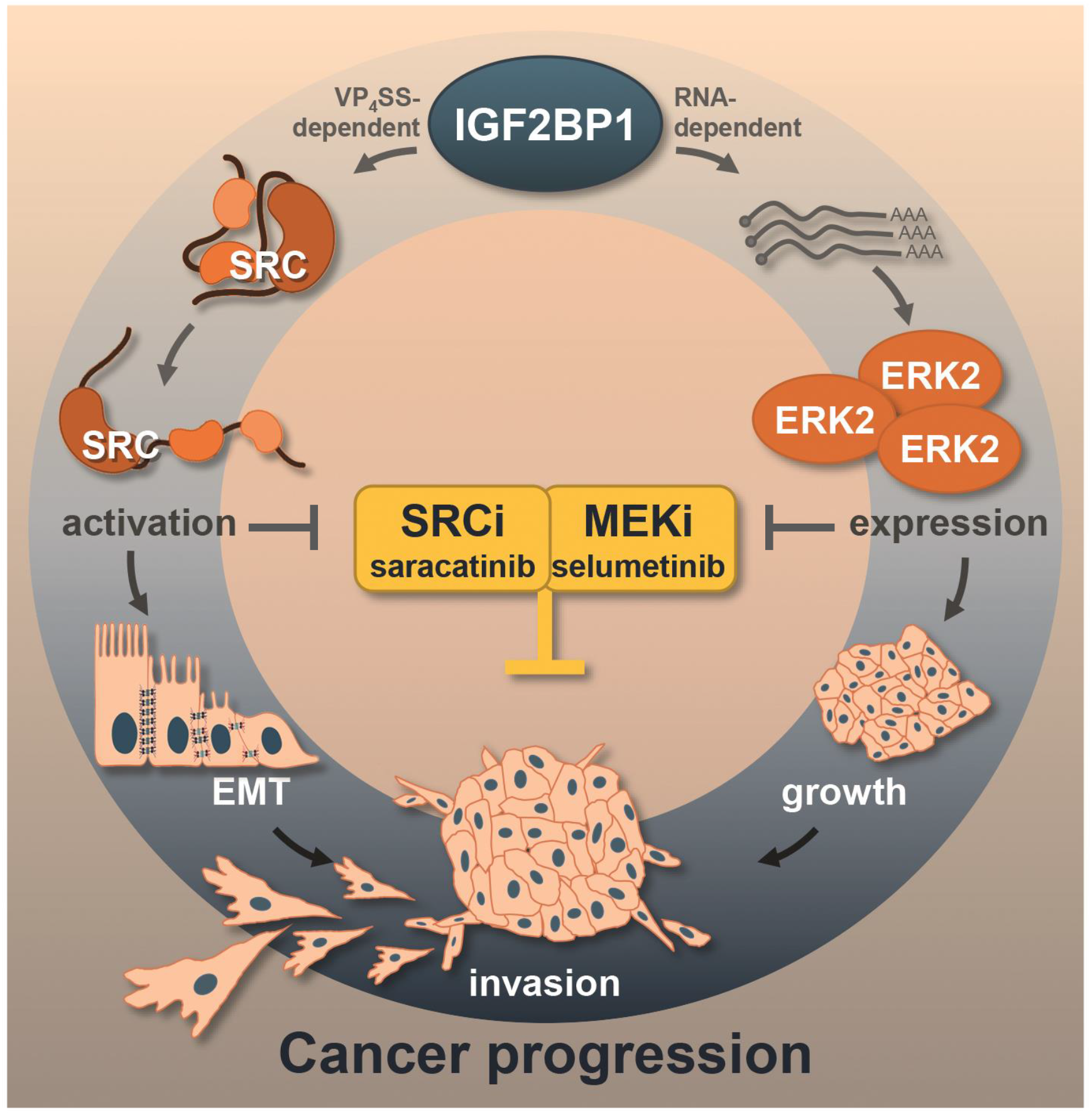

## Introduction

EOC (epithelial ovarian carcinoma) has the highest death rates among female cancers and improvement of therapeutic efficacies remains dismal [1]. Although promising initially, PARP- and SRC-directed therapies suffer from occurring resistances and the lack of biomarkers required to identify patients benefitting from the respective treatments [2–5].

Unsupervised clustering of patient samples by transcriptome data identified molecular subtypes of HG-SOCs with distinct characteristics, including an invasive, mesenchymal-like subtype, observed in approximately 30% of patients [1, 6–8]. EMT, characterized by the loss of CDH1 (E-cadherin) positive cell-cell contacts termed adherens junctions (AJs), is thought to underly progression of the mesenchymal-like subtype. The SRC protein kinase is a crucial activator of AJ disassembly with elevated activity in EOC patients [9, 10]. SRC phosphorylates AJ components like CTNNB1 (β-Catenin), CDH1 or CDH2 (N-cadherin) resulting in their internalization and/or degradation [9, 11–13]. In EOC-derived OVCAR-3 cells, SRC promotes EMT, and this activity is further pronounced by ERK2 [14]. ERK2 activates SRF (serum response factor) driving the transcription of ECM molecules and integrins [15]. This was suggested to promote tumor growth, mesenchymal properties and metastasis. Accordingly, small molecule inhibitors of SRC and ERK1/2-activating MEK, in particular saracatinib and selumetinib, are tested individually in clinical trials (ClinicalTrials.gov Identifiers: NCT01196741; NCT00610714; NCT00551070; NCT03162627) [5, 16]. In EOC xenograft models, their co-application proved beneficial over single drug therapies, since MEK inhibition re-sensitized saracatinib resistant cells [2, 10]. These findings provide preclinical evidence suggesting that the combination of SRC and MEK inhibitors is beneficial in the treatment of ovarian cancer, and highlights the need for identifying biomarkers for selecting patients for clinical trials [2].

IGF2BP1 is an oncofetal RNA-binding protein (RBP) up-regulated in a variety of solid tumors [17]. In cancer, IGF2BP?s main function is the partially m^6^A-dependent (N^6^-methyladenosine) impairment of miRNA-directed mRNA decay [15, 18–23]. This results in elevated expression of oncogenic factors like LIN28B, HMGA2, MYC, SRF and ERK2. During development, the spatially restricted synthesis of ACTB is essentially controlled by the SRC-directed phosphorylation of IGF2BP1 [24]. SRC associates with IGF2BP1 at a SH3-binding VP4SS motif conserved in the SRC substrate and cell adhesion protein Paxillin (PXN) [25].

IGF2BP1 is up-regulated in EOC and associated with adverse patient outcome [19, 26]. Its depletion or deletion decreases the spheroid growth, migration, and invasion of ovarian cancer cells *in vitro* and reduces tumor growth as well as metastasis *in vivo* [18, 19, 27]. In melanoma-derived cell lines, IGF2BP1 enforces mesenchymal tumor cell properties in a LEF1-dependent manner [28]. However, if and how IGF2BP1 modulates EMT in EOC remained elusive.

Here we reveal a novel, RNA-independent mechanism of IGF2BP1-dependent SRC activation. In concert with enhancing ERK2 expression, IGF2BP1 promotes SRC/ERK2-signaling in EOC-derived cells providing a rational for the therapeutic inhibition of SRC and MEK in mesenchymal-like HG-SOC.

## Results

### IGF2BP1 is a pro-mesenchymal driver up-regulated in the C5 subtype of HG-SOC

Consistent with previous findings, IGF2BP1 expression is associated with adverse prognosis in EOC tumors [18, 19, 26] (Supplementary Figure 1A,B). Correlation analyses in three independent transcriptome datasets indicated that IGF2BP1 mRNA expression is strongly associated with the C5 subtype of HG-SOC (Figure 1A; Supplementary Figure S1C-F). In agreement, IHC (immunohistochemistry) revealed a higher Remmele score for IGF2BP1 in C5 tumors derived from a local tumor cohort (Figure 1B,C; Supplementary Table T1B). To identify candidate effector pathways of IGF2BP1 in EOC, the TCGA-provided transcriptome data set was separated in IGF2BP1 low (< 5 cpm) and high (> 5 cpm) expressing tumors. Median IGF2BP1 mRNA expression was more than 25-fold up-regulated in one third of patients (Figure 1D; Supplementary Table T1A). Gene set enrichment analyses (GSEA) using the fold change of gene expression identified significant up-regulation of proliferation- and EMT-associated gene sets in IGF2BP1-high vs. low tumors (Figure 1E; Supplementary Figure S1G and Tables T1A-3). In agreement with a pro-mesenchymal role of IGF2BP1, the protein was markedly elevated in a subset of mesenchymal-like EOC-derived cell lines (Figure 1F,G). These were characterized by high abundance of mesenchymal markers VIM, CDH2 and ZEB1 and low levels of epithelial markers CDH1, KRT8 and EPCAM (Figure 1F,G; Supplementary Table T4). Immunostaining of CTNNB1 and F-actin labeling confirmed a pronounced mesenchymal-like morphology of ES-2 cells with diminished CTNNB1-positive cell-cell contacts when compared to epithelial-like OVCAR-3 cells (Figure 1H, I). To test if IGF2BP1 promotes a mesenchymal-like phenotype in EOC-derived cells, the protein was depleted or over-expressed in a panel of EOC-derived cells. IGF2BP1 depletion reduced 3D spheroid growth and invasion in all EOC cell lines tested (Figure 1 J,K; blue boxes). Strikingly, the forced expression of GFP-fused IGF2BP1 significantly elevated the invasive growth of OVCAR-3 and ES-2 cells (Figure 1 J,K; red boxes). In sum, this indicated that IGF2BP1 is a marker of the C5 subtype of HG-SOC, promoting invasive growth in EOC-derived cell models.

**Figure 1.**
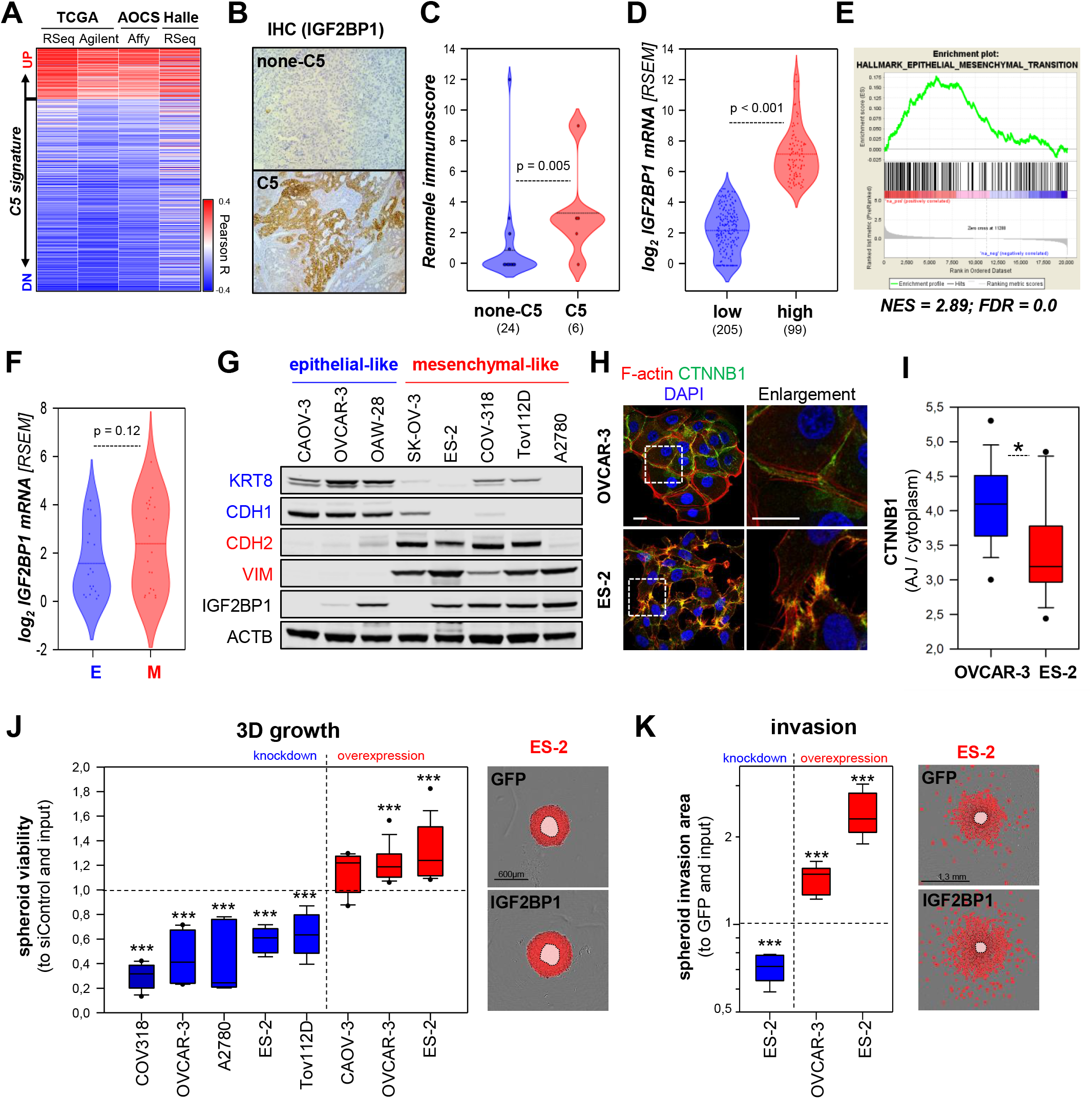
IGF2BP1 is associated to the C5 signature and promotes mesenchymal properties. **(A)** IGF2BP1 expression was correlated to the C5 signature using indicated data sets. Pearson correlation coefficients (R) are shown as heat map. **(B,C)** IHC staining of EOC samples classified as C5 or none-C5 via NGS based GSEA analyses using IGF2BP1 directed antibodies (B). IGF2BP1 staining was quantified using the Remmele immune score in none-C5 (24) and C5 (6) samples (C). **(D)** IGF2BP1 mRNA expression is shown by violin plots, in the TCGA-OV-RNA-Seq cohort distinguished in IGF2BP1-high (log_2_ RSEM ≥ 5) or -low (log_2_ RSEM ≤ 5). **(E)** GSEA plot of the HALLMARK_EMT gene set based on gene ranking by fold change expression between IGF2BP1-high vs -low samples as in (D). **(F)** Violin plot of IGF2BP1 mRNA expression in EOC-derived cells, classified as epithelial-like (E, blue) or mesenchymal-like (M, red) by the differential mRNA expression (Cancer Cell Line Encyclopedia; CCLE expression data) of epithelial (CDH1, KRT8 and/or EPCAM) and mesenchymal (VIM, ZEB1 and/or CDH2) markers. **(G)** Western blotting of IGF2BP1 and indicated epithelial (blue) and mesenchymal (red) markers. VCL (vinculin) served as loading control. **(H,I)** Immunostaining of CTNNB1 and F-actin using Phalloidin-TRITC in indicated cell lines. Dashed boxes depict enlarged regions. Scale bar 25 μm. The ratio of CTNNB1 membrane to cytoplasmic localization was determined in 15 cells over three independent analyses. **(J)** 3D growth of indicated cell lines upon IGF2BP1 knockdown (blue) or over-expression (red), as determined by Cell Titer Glo relative to controls and input. Representative images of ES-2 cells expressing GFP or GFP-IGF2BP1 overlaid with growth detection masks (red) and input spheroids (light red) are shown. **(K)** Invasion of pre-formed Matrigel embedded spheroids of ES-2 cells upon IGF2BP1 knockdown (blue) or over-expression (red) was monitored in an Incucyte S3. Representative images overlaid with invasion detection masks (red) or input spheroids (light red) are shown. Box plots in J and K are derived from analyses of 6-9 spheroids in three independent experiments. Statistical significance was determined by Student’s T-test or Mann-Whitney rank sum test. *, p < 0.05; ***, p < 0.001.

### IGF2BP1 promotes the disassembly of adherens junctions

How aberrant IGF2BP1 expression modulates cell morphology was investigated in epithelial-like OVCAR-3 and mesenchymal-like ES-2 cells. IGF2BP1 over-expression in OVCAR-3 enhanced the formation of lamellipodia and led to partial disassembly of cell colonies (Figure 2A, arrow heads). IGF2BP1 depletion in ES-2 cells reduced spindle-like morphologies in favor of a compact, flattened shape and enhanced the formation of colonies, as previously observed [28]. This implied IGF2BP1-dependent regulation of cell-cell contacts, in particular the modulation of AJ integrity, analyzed further by CDH1 and CTNNB1 immunostaining. IGF2BP1 over-expression in OVCAR-3 cells converted mature AJs to premature, zipper-like AJs (Figure 2B). Quantification of AJ/cytoplasm fluorescence intensities confirmed decreased AJ localization of CDH1 and CTNNB1, suggesting enhanced internalization and/or decay due to IGF2BP1 over-expression.

**Figure 2.**
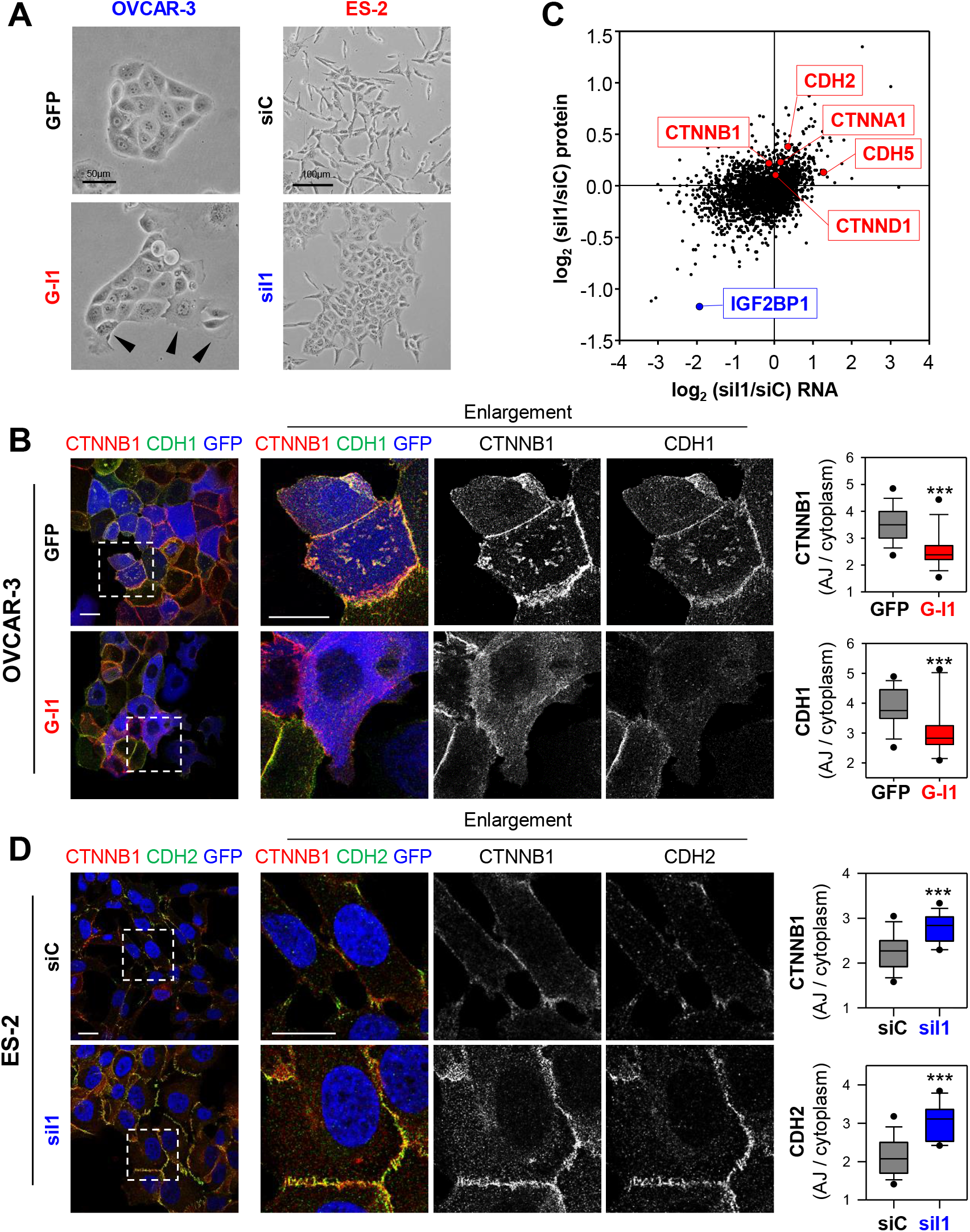
IGF2BP1 promotes AJ disassembly. **(A)** Phase contrast images of OVCAR-3 and ES-2 cells upon GFP or GFP-IGF2BP1 (G-I1) over-expression and IGF2BP1 (siI1) or control depletion (siC). Arrowheads indicate colony-detaching cells. **(B)** Immunostaining of IGF2BP1 over-expressing or control OVCAR-3 cells with indicated antibodies. Over-expression is pseudo-colored in blue. Dashed boxes depict enlarged region shown in middle panels. The localization of CDH2 and CTNNB1 was assessed by immunostaining. The membrane/cytoplasmic ratio of fluorescence intensities is shown by box plots (right panels). **(C)** Log_2_ fold change of protein expression (quantitative proteomics) upon IGF2BP1 depletion in ES-2 cells plotted over the respective log_2_ fold changes in RNA abundance (NGS). Selected AJ proteins are indicated in red, IGF2BP1 is highlighted in blue. **(D)** ES-2 cells with transient IGF2BP1 depletion were analyzed as in (B). Scale bars 25 μm. 15 cells per condition were analyzed in three independent experiments. Statistical significance was tested by student’s T-test. ***, p < 0.001.

In ES-2 cells, the depletion of IGF2BP1 induced severe morphological changes reminiscent of a partial mesenchymal-to-epithelial (MET) transition (Figure 2A and D). This was indicated by markedly increased CDH2 and CTNNB1 AJ-recruitment, although CDH1 expression was not restored (Figure 2D). Furthermore, IGF2BP1 knockdown enforced reorganization of the intermediate as well as microfilament system and induced the formation of premature AJs (Supplementary Figure S2A-C). These even showed increased recruitment of the AJ-stabilizing factor CTNND1 (δ-Catenin) (Supplementary Figure S2B; [29]). MET-like transition, indicated by enriched AJ-localization of CDH2 and CTNNB1 was conserved in a panel of EOC-derived cells and cell lines of other origin depleted for IGF2BP1 (Supplementary Figure S2D,E). The deletion of IGF2BP1 in ES-2 cells led to drastic MET-like morphological changes culminating in the formation of mature AJs (Supplementary Figure S3A). On the contrary, IGF2BP1 over-expression further reduced the AJ-localization of CTNNB1 and CDH2 in ES-2 and COV-318 cells (Supplementary Figure S3B,C). This re-localization was essentially abolished by depleting exogenous IGF2BP1 in ES-2 (Supplementary Figure S3B).

How IGF2BP1 influences the expression of AJ proteins, was analyzed further by monitoring mRNA and protein abundance by mRNA-sequencing and quantitative MS analyses upon IGF2BP1 depletion in ES-2 cells (Figure 2C; Supplementary Table T5). This was associated with up-regulation of AJ proteins CDH2, CDH5, CTNNA1 and CTNNB1. Together these observations supported the notion that IGF2BP1 up-regulation is a driver of AJ disassembly and EMT in EOC cells.

### IGF2BP1 regulates SRC activity

IGF2BP1 is a key regulator of mRNA turnover in cancer cells, suggested to control CTNNB1 mRNA turnover in breast cancer cells [15, 19, 20, 30]. However, with the exception of CDH5, steady state mRNA levels or mRNA turnover of AJ components including CTNNB1 remained unchanged upon IGF2BP1 depletion (Figure 2C; Supplementary Figure S4A). Moreover, the activity of CDH2- and CTNNB1-3’UTR luciferase reporters remained unaffected by IGF2BP1 knockdown (Supplementary Figure S4B). This largely excluded 3’UTR-dependent regulation by IGF2BP1, as previously reported for ACTB and MAPK4 mRNAs [24, 27]. Instead, our findings implied that elevated AJ formation upon IGF2BP1 depletion, involves altered turnover of AJ proteins [29, 31]. In agreement, the decay of CDH2 and CTNNB1 proteins was significantly reduced by IGF2BP1 knockdown (Figure 3A,B).

**Figure 3.**
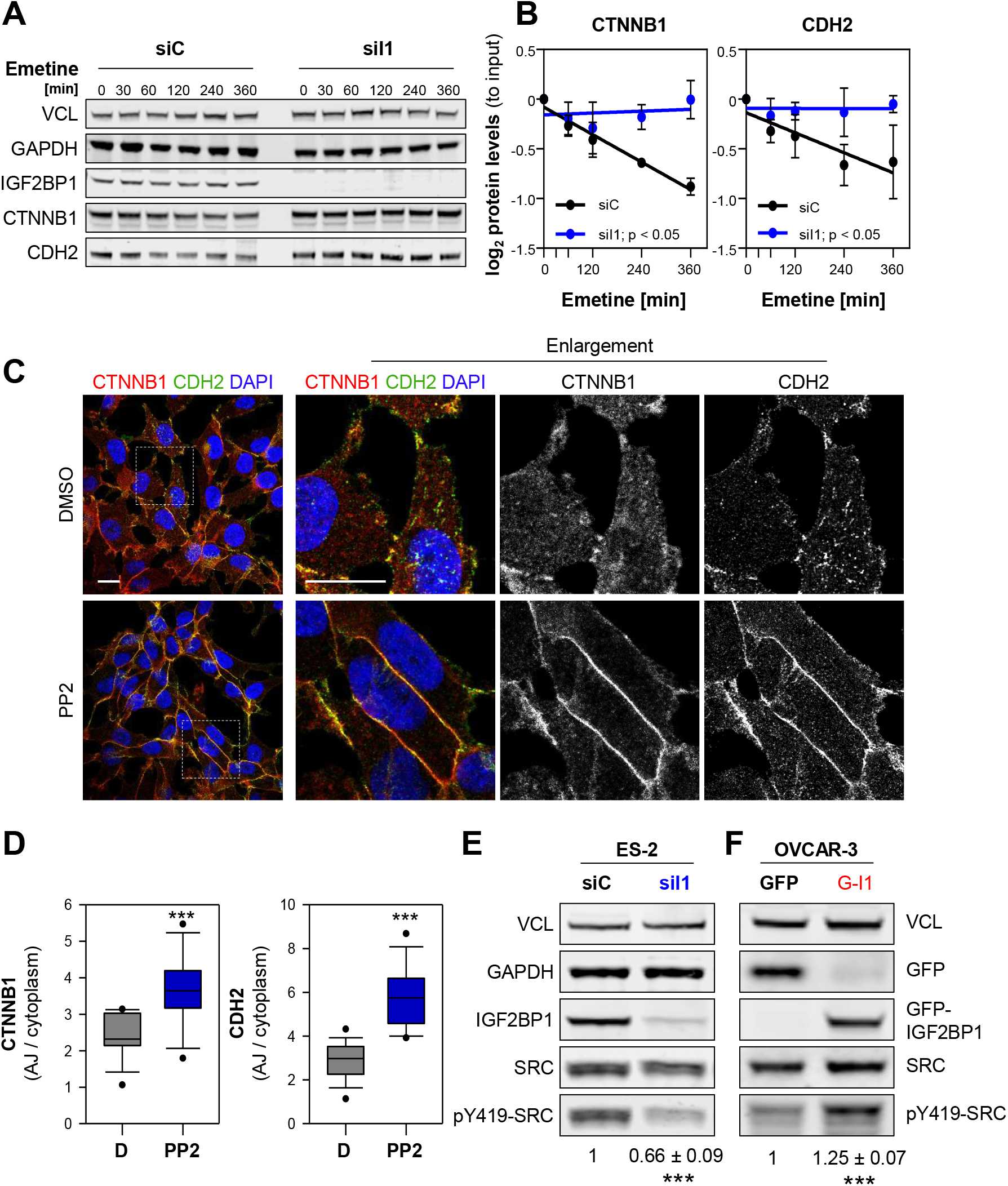
IGF2BP1 destabilizes AJs in a SRC-dependent manner. **(A,B)** Turnover of indicated proteins was determined by blocking protein synthesis with Emetine (100μM) by Western blotting upon IGF2BP1 (siI1) and control (siC) knockdown. GAPDH and VCL served as loading controls to determine protein abundance relative to input levels. **(C,D)** Immunostainings of ES-2 cells treated with PP2 (1μM) or DMSO for 48h using indicated antibodies (C). Dashed box indicates enlargement. Scale bars 25 μm. The AJ/cytoplasmic ratio of CDH2 and CTNNB1 is shown by box plots (D). **(E,F)** Western blot analyses of indicated proteins in ES-2 cells upon IGF2BP1 depletion (E) and OVCAR-3 cells (F) with GFP or GFP-IGF2BP1 (G-I1) over-expression. SRC activity, indicated by Y419 phosphorylation, was normalized to SRC expression, as depicted under the lanes. VCL served as loading control. Error bars show SD determined in three independent experiments. Statistical significance was tested by Student’s T-test. *, p < 0.05; ***, p < 0.001.

The phosphorylation of some AJ proteins, e.g. CDH2, by SRC kinase promotes AJ disassembly and subsequently protein decay [9, 11–13]. Consistently, AJ localization and total abundance of CTNNB1 and CDH2 were elevated upon treatment of ES-2 cells with the SRCi PP2 (Figure 3C,D; Supplementary Figure S4C). In view of IGF2BP1’s association with the SH3 domain of SRC [24], it appeared tempting to speculate that IGF2BP1 influences SRC activity. Western blotting revealed that SRC activity, indicated by phosphorylation at tyrosine 419 (Y419) in the kinase domain, was substantially reduced upon IGF2BP1 depletion in all EOC-derived cell lines tested (Figure 3E; Supplementary Figure S4D). Conversely, increased Y419-SRC phosphorylation was observed upon the over-expression of IGF2BP1 in OVCAR-3 cells (Figure 3F). SRC expression remained essentially unchanged by aberrant IGF2BP1 expression. In sum, these findings suggested that IGF2BP1-directed activation of SRC promotes AJ disassembly in EOC-derived cells.

### IGF2BP1 promotes SRC activation via a SH3-ligand-binding mechanism

Previous studies identified IGF2BP1 as a SRC substrate and suggested association of the kinase’s SH3 domain at a VP4SS motif in IGF2BP1 [24]. This putative SRC-binding site is highly conserved among IGF2BP1 orthologues and identical to the SRC-docking site of the adhesion molecule Paxillin (Figure 4A) [24, 25, 32]. SRC activity is repressed by an intramolecular interaction (closed conformation) [33]. Activation, indicated by an open conformation enabling auto-phosphorylation at pY419 in the kinase domain, is controlled by various means including ligand-binding induced mechanisms [34]. The latter were described for some SRC substrates including Paxillin and the RNA-binding proteins hnRNPK and SAM68 [32, 35, 36]. To test if IGF2BP1 promotes SRC activity via its VP4SS motif, increasing amounts of GFP-tagged proteins were expressed in ES-2 cells and SRC activity was monitored by Western blotting for pY419-SRC (Figure 4B,C). This revealed dose-dependent up-regulation of pY419-SRC by wild type (WT) and RNA-binding deficient (KH) IGF2BP1 (Figure 4B,C) [19, 37]. In contrast, SRC activity remained unchanged upon deletion of the VP4SS motif in IGF2BP1 (ΔSH3). Although less efficient, SRC-activity was also enhanced by over-expression of the GFP-fused VP4SS peptide (SH3motif). In sum, this unraveled that IGF2BP1 promoted SRC activity in a dose-dependent manner by a ligand-binding induced, but RNA-binding independent mechanism.

**Figure 4.**
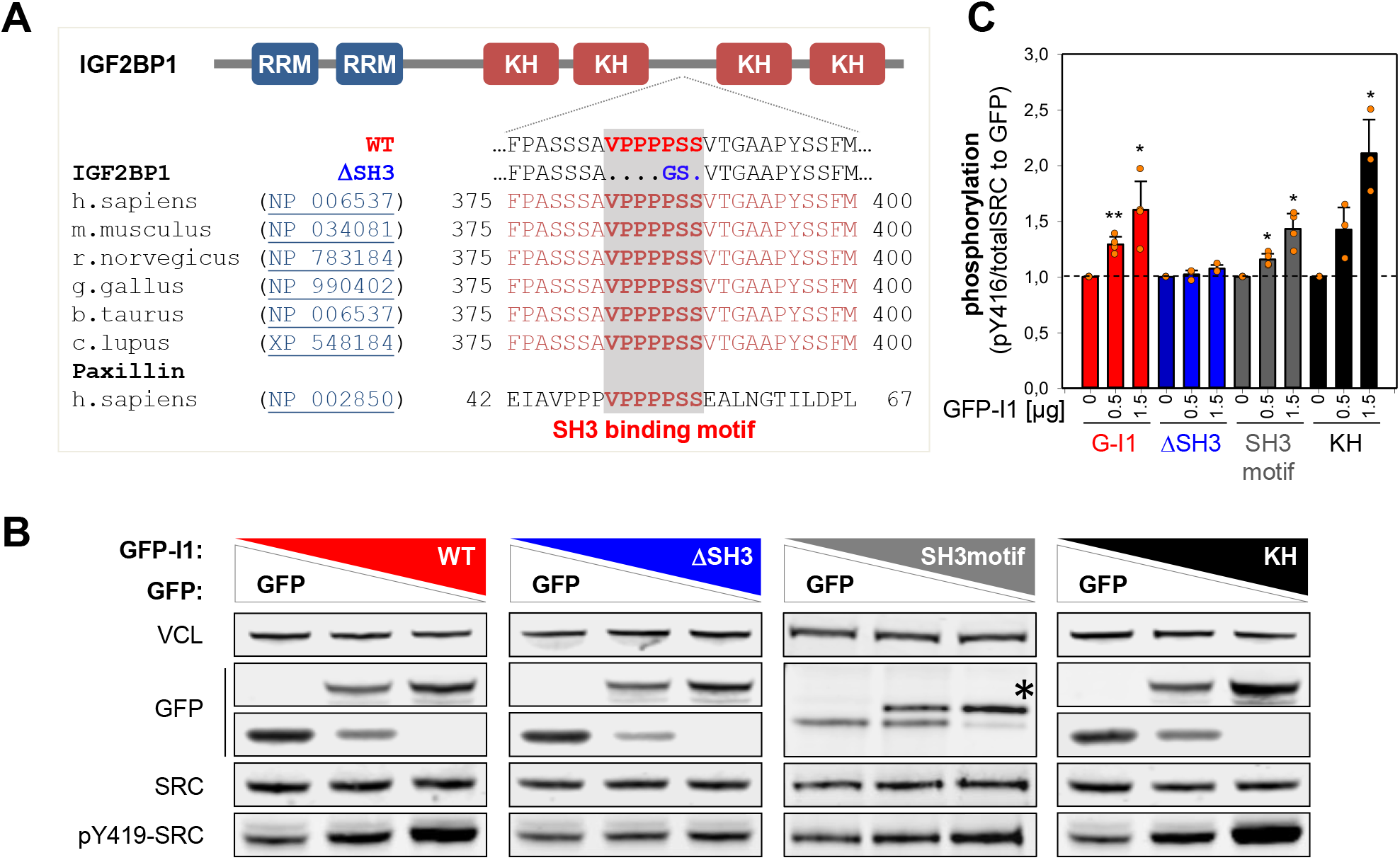
IGF2BP1 promotes SRC activation via its SH3-binding VP4SS motif. **(A)** Schematic representation of VP4SS conservation among IGF2BP1 orthologs and human paxillin. The domain organization of IGF2BP1 is shown in the upper panel (RRM, RNA recognition motif; KH, hnRNPK homology domain). **(B)** Representative Western blot analyses of indicated proteins in ES-2 cells transiently expressing GFP alone (lane 1) a mixture of GFP and indicated GFP-fused proteins (lane 2) or only GFP-fused proteins. WT, GFP-IGF2BP1WT; DSH3, GFP-IGF2BP1DSH3; SH3motif, GFP-VP4SS; KH, GFP-IGF2BP1KHmut. VCL served as loading control. **(C)** Quantification of Western blots shown in (B). Error bars indicate SD determined in three independent experiments. Statistical significance was tested by Student’s T-test. *, p < 0.05; **, p < 0.01.

### The IGF2BP1-SRC axis enhances EMT and invasive growth of EOC-derived cells

The physiological relevance of IGF2BP1-dependent control of SRC activity was analyzed by monitoring AJ integrity using immunostaining (Figure 5A,B). IGF2BP1 over-expression in OVCAR-3 or re-expression in IGF2BP1-deleted ES-2 cells impaired the AJ localization of CTNNB1, CDH1 or CDH2 compared to GFP controls. Similar to wild type IGF2BP1, the RNA-binding deficient (KH) protein substantially impaired AJ assembly. In contrast, AJ localization of the respective proteins was essentially unchanged by over- or reexpression of IGF2BP1 lacking the VP4SS motif (ΔSH3).

**Figure 5.**
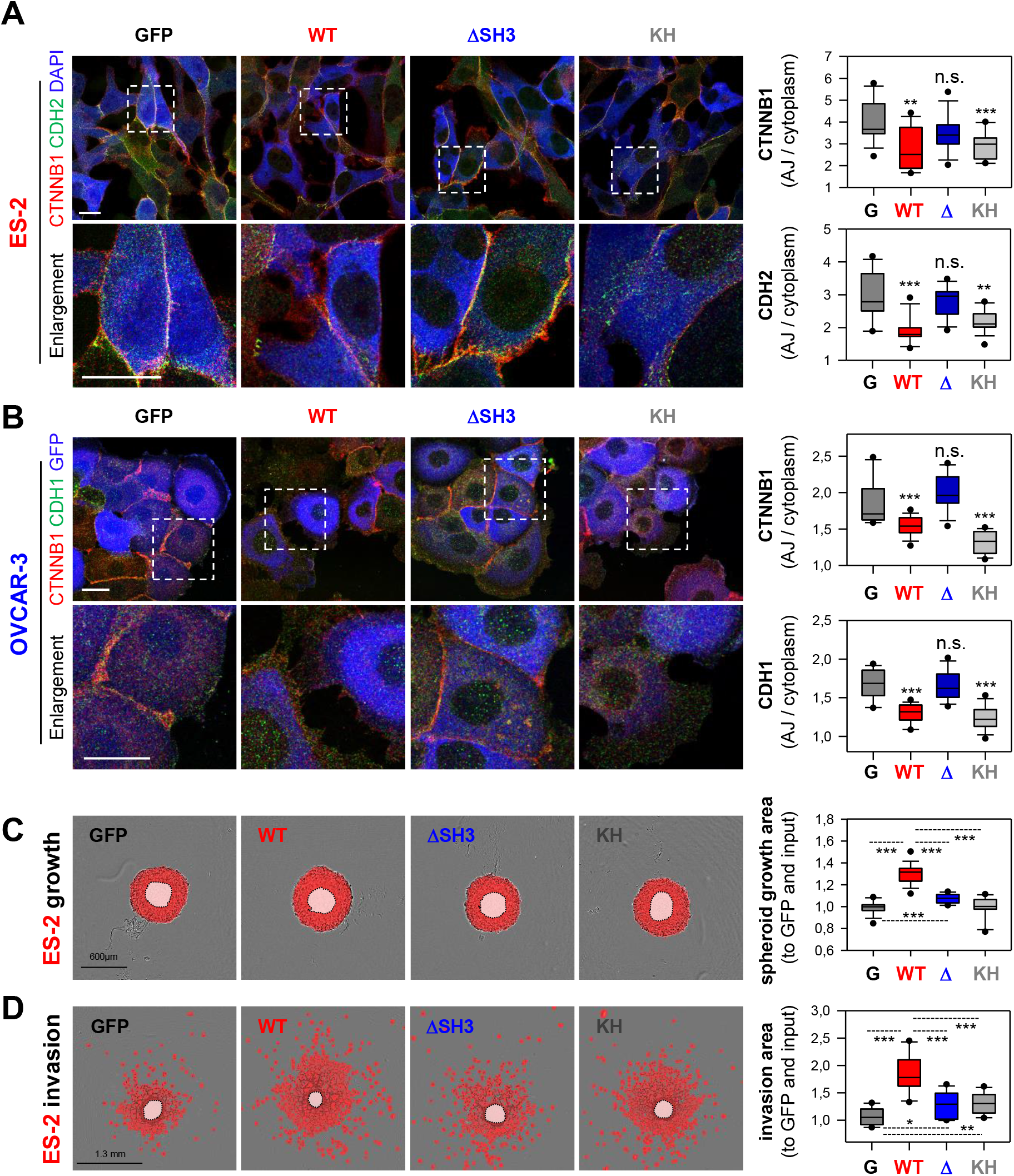
IGF2BP1 promotes AJ disassembly VP_4_SS-dependent. **(A,B)** Immunostaining of CTNNB1 and CDH2 or CDH1 upon the over-expression of GFP or indicated IGF2BP1 proteins (pseudo-colored in blue) in IGF2BP1-deleted ES-2 (A) or parental OVCAR-3 cells (B) were performed and analyzed as in Figure 1H,I. Dashed boxes indicate enlargements. Scale bars 25 μm. **(C,D)** Spheroid growth (C) or invasion (D) of ES-2 cells stably expressing GFP or indicated IGF2BP1 proteins were performed and analyzed as in Figure 1J,K. Statistical significance was tested by Student’s T-test or Mann-Whitney rank sum test. *, p < 0.05; **, p < 0.01; ***, p < 0.001.

However, only wild type IGF2BP1 substantially promoted both, 3D spheroid growth and invasion of ES-2 cells (Figure 5C,D). In contrast, invasive growth was only modestly elevated by ΔSH3 and KH mutant proteins. To investigate the role of IGF2BP1-dependent SRC-activation in invasive growth, the kinase was inhibited by PP2 in ES-2 cells over-expressing IGF2BP1 (Supplementary Figure S4E). This essentially abolished IGF2BP1-directed invasion. Collectively, these findings indicated a pivotal role of IGF2BP1-directed SRC activation in AJ integrity and invasive growth of EOC-derived cells. However, full potential in promoting invasive growth requires both, the RNA-independent activation of SRC and presumably the RNA-dependent regulation of mRNA fate by IGF2BP1.

### Combined SRCi/MEKi effectively impairs IGF2BP1-driven invasive growth

The SRC inhibitor saracatinib showed anti-tumor activity in HG-SOC preclinical studies [38]. However, activation of the MAPK pathway is associated with resistance to saracatinib. MEK inhibition by selumetinib proved effective in re-sensitizing saracatinib resistant cells [2]. Co-activation of SRC and MAPK was reported in HG-SOC patients and combined SRCi/MEKi impaired tumor growth *in vivo* more effectively than monotherapies [10]. We recently demonstrated that IGF2BP1 promotes ERK2 (MAPK1) and SRF expression by impairing miRNA-directed repression [15, 19]. Consequently, SRF-dependent MAPK signaling was reduced upon IGF2BP1 depletion [15]. Analyses in a panel of EOC-derived cells confirmed, IGF2BP1 is a conserved enhancer of ERK2 expression and MAPK signaling (Supplementary Figure S5A-C).

Currently, no clinically evaluated IGF2BP1 inhibitor is available. The small molecule inhibitor BTYNB impairs tumor cell growth and IGF2BP1-RNA binding *in vitro* [39]. However, it appeared unlikely to inhibit the RNA-binding independent control of SRC activity by IGF2BP1. Hence, we analyzed if saracatinib and/or selumetinib impair IGF2BP1-directed control of invasive growth.

To this end, we determined the EC_50_ concentrations for both inhibitors in the 3D growth and invasion of ES-2 cells over-expressing GFP or GFP-IGF2BP1. Irrespective of IGF2BP1 over-expression, selumetinib repressed 3D growth and saracatinib invasion at nearly equal potency (Supplementary Figure S5D,E). Conversely, IGF2BP1 over-expression induced a 60% higher tolerance towards saracatinib under 3D growth conditions and four-fold higher invasion upon selumetinib treatment (Figure 6A,B). In support of this, EOC-derived cell lines were sensitized to growth inhibition by saracatinib (Sa; approximated EC_50_ concentration of 3μM) upon IGF2BP1 depletion and became more resistant by IGF2BP1 over-expression (Figure 6C). The inhibition of invasion by selumetinib (Se; approximated I_max_ concentration of 3μM) was elevated by IGF2BP1 depletion and decreased by its over-expression (Figure 6D).

**Figure 6.**
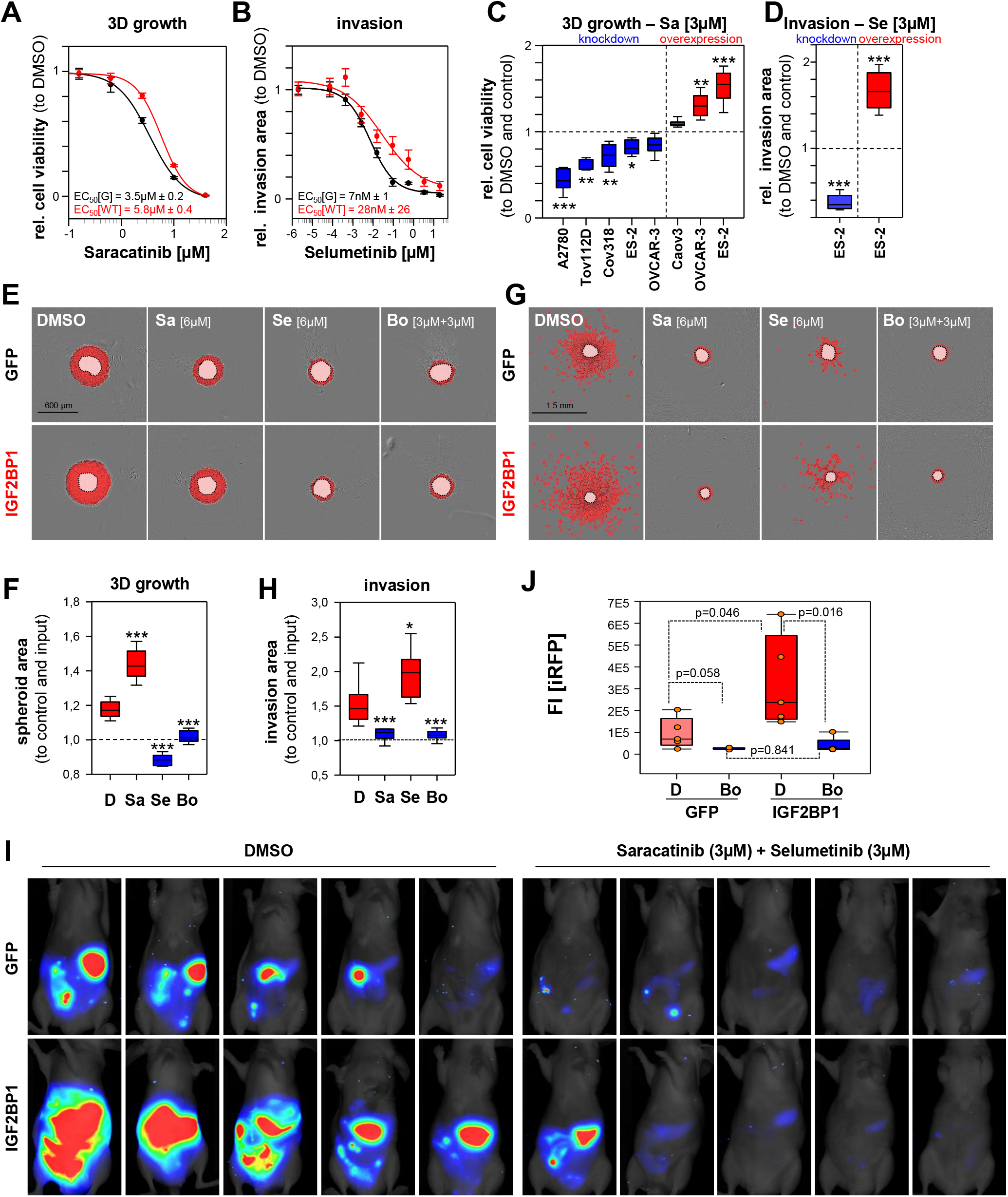
IGF2BP1 promoted invasive growth is abolished by combined SRCi/MEKi treatment. **(A-H)** Spheroid growth (A,C,E,F) or invasion (B,D,G,H) of GFP (controls) or IGF2BP1 over-expressing ES-2 cells was monitored and analyzed as in Figure (1J,K). All treatments at indicated concentrations were performed on pre-formed (24h) spheroids for indicated time and concentrations. Error bars indicate SE (A,B). **(I,J)** iRFP-labelled GFP or IGF2BP1 over-expressing ES-2 cells pre-treated with saracatinib (3μM) and selumetinib (3μM) or DMSO for 48h, were IP injected into nude mice with compounds at indicated concentrations. Tumor growth of five mice per conditions (I) was monitored and quantified (J) by infra-red imaging of the abdominal region 2 weeks post injection. Statistical significance was tested by Student’s T-test or Mann-Whitney rank sum test. *, p < 0.05; **, p < 0.01; ***, p < 0.001.

To address if both inhibitors act in a synergistic or additive manner, combinatorial treatment was performed using a 2D growth drug matrix analysis (Supplementary Figure S6A, B). This revealed additivity, indicated by synergy scores in-between −10 and 10, determined by three independent synergy models [40]. The maximal additive concentrations for IGF2BP1-expressing were approximately 6μM for both inhibitors. According to HSA model assumptions [40], the potential benefit of additive inhibition in 3D growth and invasion was analyzed at 3μM per drug in combined and 6μM in mono-treatments. Western blotting confirmed inhibition of both kinases by the respective inhibitors at these concentrations (Supplementary Figure S6C).

However, even at 6μM single saracatinib treatment remained substantially less efficient than selumetinib or combined treatment in 3D growth analyses (Figure 6E,F; Supplementary Figure S6D). The same was observed for selumetinib in 3D invasion studies (Figure 6G, H; Supplementary Figure S6E). Thus, only combined treatment was effective in inhibiting both, 3D growth and invasion irrespective of IGF2BP1 over-expression.

Invasive growth in the peritoneum is a major complication in the progression of ovarian cancer. If combined treatment also impairs invasive growth *in vivo* was analyzed by intraperitoneal (IP) injection of iRFP-labeled (near-infrared red fluorescent protein) ES-2 cells expressing GFP or GFP-IGF2BP1. Cells were pre-treated with both inhibitors (3 μM each), and DMSO for 48h and injected in the presence of both inhibitors at the same concentrations. Tumor growth was monitored by total iRFP signal in the peritoneum two weeks post injection of viable tumor cells (Figure 6I,J). IGF2BP1 over-expression led to higher tumor burden in DMSO treated controls, supporting the growth- and invasion-enhancing role of IGF2BP1 observed *in cellulo* and previously reported [19]. Irrespective of IGF2BP1 expression, tumor growth was essentially abolished by combined treatment.

Together these observations indicated that combined treatment with selumetinib and saracatinib effectively impairs the IGF2BP1-dependent enhancement of invasive growth in EOC-derived cells *in vitro* and *in vivo*.

## Discussion

Enhanced expression of IGF2BP1 is associated with a mesenchymal/proliferative-like gene signature observed in approximately 30% of serous ovarian cancers [6–8]. This identifies IGF2BP1 as a novel marker of the mesenchymal-like C5 subtype of HG-SOCs [6]. In EOC-derived cells, IGF2BP1 promotes invasive growth by stimulating SRC/ERK-signaling. The protein enhances ERK2 abundance by stabilizing the respective mRNA, as previously reported [19]. In contrast, SRC-activation by IGF2BP1 is RNA-independent and solely relies on the conserved SRC/SH3-binding motif of IGF2BP1. This suggests that IGF2BP1 promotes SRC kinase activation by a ligand-binding-induced mechanism prior unknown for IGF2BP1. This is highly important for perspective drug-design approaches currently addressing IGF2BP1’s RNA binding activity only [39]. However, full potential of IGF2BP1 enhanced invasive growth requires both, protein-dependent activation of SRC and RNA-binding associated regulation.

The activation of SRC is crucial for the pro-invasive role of IGF2BP1, since it promotes EMT by inducing the disassembly of AJs. This is abolished by ablating SRC activity using pharmacological inhibitors. This reveals a fundamental novel role of IGF2BP1, the interconnection of deregulated gene expression and cancer cell signaling. Consistent with this dual function promoting ERK/SRC signaling, the combined inhibition of MAPK- and SRC-signaling by selumetinib and saracatinib proves substantially more effective than monotherapies. This was validated IGF2BP1-dependent for the impairment of 3D growth and invasion *in cellulo* and the peritoneal spread of tumor cells in experimental mouse tumor models. In support of this, previous studies report a substantial therapeutic benefit of combined inhibition of MAPK/SRC signaling in EOC tumor models [2, 10]. Thus, our findings suggest IGF2BP1 as a novel marker for EOC therapy. Considering the major problem of invasive growth associated with ovarian cancer spread in the abdominal cavity, the further investigation of IGF2BP1-driven cancer progression, in particular its roles in the mesenchymal subtype of HG-SOCs, appears mandatory. Along these lines, the direct targeting of IGF2BP1-dependent functions, for instance by impairing its RNA-binding by small molecules like BTYNB need to be evaluated.

## Materials and methods

### Patient samples and IHC

Human HG-SOC samples, collected from patients undergoing surgery, were prepared by the pathologist and immediately frozen (liquid nitrogen). HE staining confirmed tumor content of each sample. The study, including IHC and NGS analyses, was approved by the Ethics Committee of the Medical Faculty/Martin Luther University Halle/Wittenberg and written informed consent. IHC staining of FFPE samples was performed as previously described [41], using a Tris/EDTA buffer (pH 9) for antigen retrieval, a commercial IGF2BP1-directed antibody (MBL, Woburn, MA;; Cat#. RN001M; RRID: AB_1953026; dilution 1:100), DAB Enhancer (Dako/Agilent, Santa Clara, CA; Cat# S1961) for detection, and counterstaining by hemalm. IGF2BP1 staining was determined blinded by two pathologists using the immunoreactive score as described [42]: 0 – absent; 1-4 – weak; 5-8 – moderate; 9-12 – strong expression. A summary of samples, IGF2BP1 expression values and Remmele scores is provided as Supplementary Table T1B.

### Animal Handling and Ethics Approvals

Immunodeficient athymic nude mice (FOXN1nu/nu) were obtained from Charles River (Wilmington, MA). Animals were handled according to the guidelines of the Martin Luther University based on ARRIVE guidelines. Permission was granted by the County Administration Office for Animal Care Saxony-Anhalt. iRFP-labelled IGF2BP1 over-expressing or GFP control ES-2 cells (7.5 x 10^4^ living cells determined by Trypan Blue counting) pre-treated with 3μM saracatinib and 3μM selumetinib or DMSO for 48h were injected intraperitoneal (IP) into six-week old female nude mice (five mice per condition) together with these inhibitors at the same concentration. Tumor growth of isofluran-anaesthetized mice was monitored weekly by near-infrared imaging using a Pearl imager (LI-COR, Lincoln, NE) and quantified using the Image Studio software (LI-COR). Mice were sacrificed after 2 weeks, as IGF2BP1 over-expressing tumors reached termination criteria.

### NGS and quantitative proteomics

NGS analyses of patient samples and cells were performed as previously described [19]. Total RNA-Seq data were deposited at NCBI GEO: (GSE147980) for EOC cohort or (GSE109605) for IGF2BP1 depletion in ES-2 cells. Quantitative proteomics were performed as described before [43]. In brief, TMT labeling of trypsin-digested proteins was performed according to the manufacturer’s instructions (TMT-10plex kit, Thermo Fisher, Waltham, MA). Equal volumes of all samples were mixed, concentrated in a vacuum concentrator and acidified with TFA for LC/MS/MS analysis. Samples were separated by nano-HPLC (Ultimate RSLC 3000, Thermo Fisher) using reversed phase C18 columns and 420-min gradients. The eluate was directly introduced into an Orbitrap Fusion Tribrid mass spectrometer (Thermo Fisher) equipped with nano-ESI source. Samples were analyzed using a collision-induced dissociation (combined ion trap-CID/high resolution-HCD) MS/MS strategy for peptide identification and reporter ion quantification. MS/MS data were searched against the NCBI database (version 140412, taxonomy homo sapiens, 89,649 entries) using the Proteome Discoverer (version 2.0, Thermo Fisher). Quantification was performed based on the reporter ion ratios derived from high-resolution MS/MS spectra. Proteomics data are shown as Supplementary Table T5.

### Public data, GSEA and databases

TCGA-OV-RNA-Seq data of the ovarian cancer cohort were obtained from GDC portal (https://portal.gdc.cancer.gov/). NGS data of ovarian cancer cell lines were downloaded from CCLE (Cancer Cell Line Encyclopedia) (https://portals.broadinstitute.org/ccle). Gene set enrichment analyses (GSEA; Supplementary Table T3) were performed as described before using the fold change expression of all genes between IGF2BP1 high and low expressing tumors (Supplementary Table T2) for gene ranking (18). Kaplan Meier plots for progression free survival (A) or overall survival (B) were generated with www.kmplot.com (3) using the microarray based cohorts for ovarian cancer including the Australian data set by Tothill et al. (4). Synergy between saracatinib and selumetinib was tested using https://synergyfinder.fimm.fi/ based on a drug matrix screen for various combinations and concentrations according to the online instructions (5).

### Cell lines, culture conditions and transfections

Ovarian cancer cell lines (Supplementary Table T6) and HEK293T cells for production of lentiviral particles were cultured in DMEM (Thermo Fisher) supplemented with 10% fetal bovine serum (FBS; Biochrom/Merck, Schaffhausen, Switzerland) at 37°C and 5% CO_2_. The most frequently used ES-2 cell line was authenticated by Eurofins Genomics using AmpFlSTR^®^ Identifiler^®^ Plus PCR Amplification Kit (Thermo Fisher) for STR profiling. Mycoplasma testing was routinely performed every two months by PCR and DAPI staining. Reverse transfections of plasmids (Supplementary Table T8) or siRNA pools (15 nM; Supplementary Table T7) were carried out using Lipofectamine 2000 or Lipofectamine RNAiMax (Thermo Fisher), respectively, according to the manufacturer’s instructions. Lenti-viral particles were produced as previously described [28].

The following inhibitors were added in indicated concentrations and time: Saracatinib (Selleckchem, Housten, TX; Cat#S1006), Selumetinib (Selleckchem; Cat#S1008) and PP2 (Sigma Aldrich, St. Louis, MO; Cat#P0042). Protein turnover was analyzed by addition of Emetine (100 μM, Sigma Aldrich, Cat# E2375) and RNA decay by addition of Actinomycin D (5μM, Sigma Aldrich, Cat# A9415) for indicated time points 72h post siRNA transfections. Protein and mRNA abundances were analyzed by Western blotting or RT-qPCR as described in the Supplementary Information. For antibodies and primers see Supplementary Table T7 and T8.

### Immunostainings

Immunostainings were essentially performed as previously described (1,2). Primary and secondary antibodies are summarized in Supplementary Table T7. Images were acquired on Leica SP5X or SP8X confocal microscopes equipped with a white light laser and HyD detectors using a 63x Oil objective and standardized settings for sequential image acquisition at the Core Facility Imaging, University of Halle. Bright field images were taken using a Nikon TE-2000-E microscope with 20x magnification and phase contrast. Representative images are shown. The localization of CTNNB1, CDH1 and CDH2 at the site of adherents junctions was quantified with respect to the cytoplasm for 14-15 cells from three independent experiments using imageJ as depicted as box plot.

### 3D spheroid growth and invasion assays

For spheroid growth or invasion assays, 1×10^3^ cells in 50 μL growth medium were seeded into round bottom ultra-low attachment plates (Corning, Corning, NY; Cat#7007), centrifuged for 5 min at 300 g and grown for 24h to induce spheroid formation. For all cell lines except ES-2 and SK-OV-3, Cultrex^®^ 10X Spheroid Formation ECM (Trevigen, Gaithersburg, MD; Cat# 3500-096-01) was added before centrifugation. For ES-2 and SK-OV-3 the addition of SFE was tested before the experiments with no improvement on 3D growth or invasion. For invasion assays, 50 μL Matrigel (Corning, Cat# 354234) was added to preformed spheroids. Growth medium (100 μL) supplemented with indicated inhibitors where stated was added before 3D growth or invasion were monitored using an Incucyte S3 (Sartorius, Göttingen, Germany) device for indicated time. Image analyses were performed using the Incucyte software for spheroid segmentation (indicated as mask overlays in the respective figures). Growth or invasion areas excluding the spheroid body are normalized to spheroid inputs. Representative images are shown. Quantifications represent in total 6-9 spheroids from three independent experiments shown as box plots.

### Antibodies, Western blotting and qRT-PCR

For Western blotting, cells were harvested by a rubber policeman to minimize degradation of trans-membrane proteins. Total protein was extracted in TLB-buffer [50 mM Tris pH 7.4, 50 mM NaCl, 1 % SDS, 2 mM MgCl_2_] followed by DNA digestion using Benzonase (Merck). Equal amounts of total protein was loaded on NuPAGE Bis-Tris (4-12%) gels (Thermo Fisher), blotted onto nitrocellulose membrane (Amersham Bioscience) using the Mini Gel Tank Blotting system (Thermo Fisher) and analyzed using the Odyssey infrared scanner (LI-COR) as previously described (26). Primary and secondary antibodies are summarized in Supplementary Table T7.

Total RNA extracted using Trizol (Thermo Fisher) served as template for cDNA synthesis by random priming using M-MLV reverse transcription system (Promega, Madison, WI). qPCR was performed based on SYBRgreen I technology using the ORA qPCR Green ROX L Mix (HighQu, Kraichtal, Germany) on a LightCycler 480 II 384 format system (Roche, Basel, Switzerland). For all primer pairs an annealing temperature of 60°C in a 3-step protocol was used. Relative changes of RNA abundance were determined by the ΔΔCt method using RPLP0 and GAPDH for normalization, as previously described (17). For primers see Supplementary Table T8.

### Plasmids, cloning and luciferase reporter assays

Plasmid generation including the respective templates, vectors, restriction sites and oligonucleotide sequences and cloning strategies are summarized in Supplementary Table T8. All PCR (polymerase chain reaction) amplified inserts were sequenced before subcloning in the respective vectors. ES-2 cells transfected with siRNAs (24h) were splitted for re-transfection of pmir-Glo luciferase plasmids (Supplementary Table T7) for additional 48h. Luciferase reporter analyses were performed essentially as previously described (17). Reporters containing a minimal vector-encoded 3’UTR (MCS) served as normalization controls.

### Statistical analyses

All cell culture experiments were performed in biological triplicates unless otherwise stated. Western blot quantifications are shown as mean ± SD. Whisker caps of box plots depict the 5^th^/95^th^ percentile and all outliers are shown. Spheroid growth and invasion curves are shown as mean ± SE. Statistical significance was tested by Student’s T-test (two-tailed) or Mann-Whitney Rank-Sum test after data distribution was checked by Shapiro-Wilk normality test using Sigma Plot.

## Acknowledgements and Funding

The work was supported by Wilhelm-Roux funding to NB and DFG funding (GRK1591) to SH. Imaging, image analyses and NGS analyses were performed at the Core Facility Imaging (CFI) of the Martin-Luther University Halle-Wittenberg, Germany.

## Disclosure of Potential Conflicts of Interests

the authors declare no conflicts of interest.

## Notes

### Competing Interest Statement

The authors have declared no competing interest.

