## Supplementary Information for "IGF2BP1 is a targetable SRC/MAPK-dependent driver of invasive growth in ovarian cancer"

**Table of content – supplementary information**

**Supplementary Figures and Legends**


| | |
|---|---|
| Figure S1 | IGF2BP1 correlates to the C5 signature of EOC tumors |
| Figure S2 | IGF2BP1 knockdown promotes AJ assembly |
| Figure S3 | IGF2BP1 promotes AJ disassembly |
| Figure S4 | IGF2BP1 controls SRC activity |
| Figure S5 | IGF2BP1 controls ERK2 expression |
| Figure S6 | Additive effects of saracatinib and selumetinib |
| Figure S7 | Full scans of Western blots for peer review |

**Supplementary Tables (excel file)**

| | |
|---|---|
| Table T1A | IGF2BP1 mRNA expression in TCGA-OV-RNA-Seq cohort |
| Table T1B | Local HG-SOC tumor cohort |
| Table T2 | differential gene expression of IGF2BP1 high vs low expressing tumors |
| Table T3 | GSEA for positive enriched gene sets |
| Table T4 | classification of EOC cell lines |
| Table T5 | quantitative proteomics |
| Table T6 | cell lines |
| Table T7 | antibodies |
| Table T8 | plasmids and oligonucleotides |

**Supplementary Table Legends**

**Supplementary Video Legends**

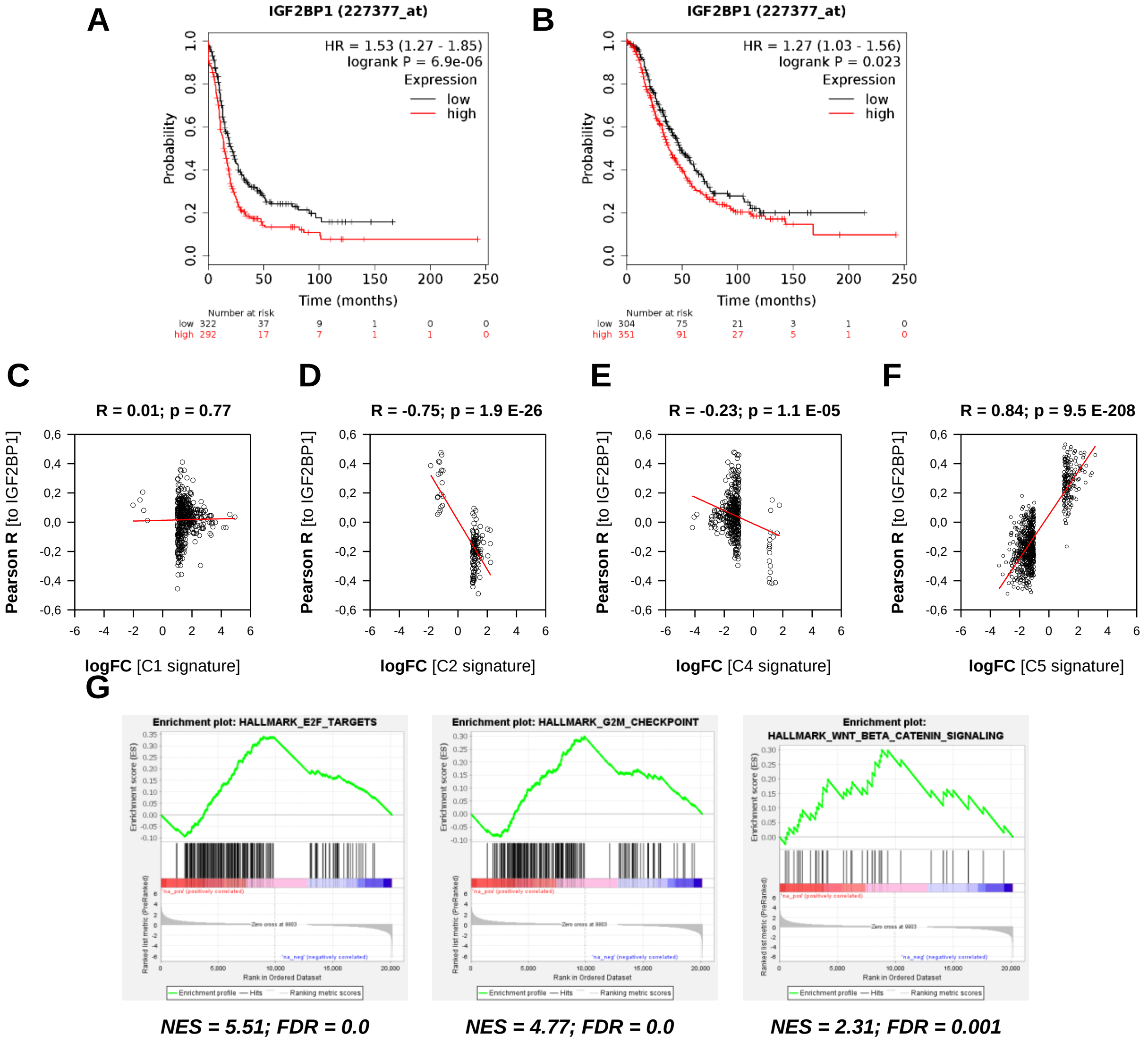


**Supplementary Figure S1. IGF2BP1 correlates to the C5 signature of EOC tumors. (A,B)** Kaplan Meier plots with indicated sample numbers were generated by www.kmplot.com using the ovarian cancer microarray based platform without any sub-selection for progression free survival (A) or overall survival (B). HR, hazardous ratio. Sample numbers for each population are indicated as numbers. **(C-F)** Pearson correlation of indicated signatures to IGF2BP1 (TCGA-OV-RNA-Seq dataset) were plotted over their respective log2 fold change determined by the signature (4). A significantly positive Pearson correlation coefficient (R) and an increasing trend line (red) indicate association of IGF2BP1 with a specific signature. This is only seen in for the C5 signature (F). **(G)** Gene set enrichment analyses (GSEA) of genes ranked by differential expression of IGF2BP1-high and low expressing tumors (TCGA-OV-RNA-Seq dataset). Please also refer to Figures 1D, 1E, and Supplementary Tables T1- T3.

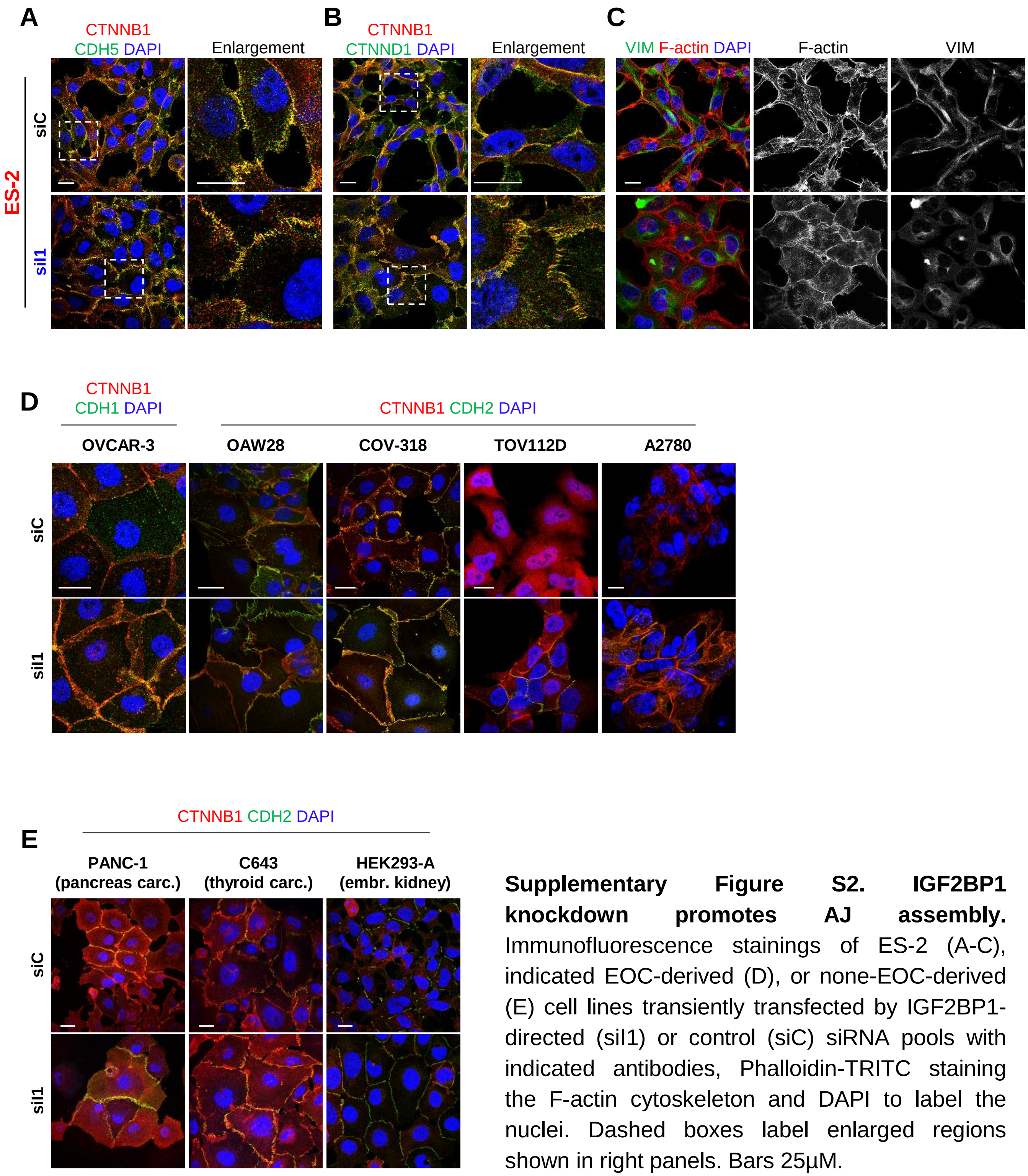


**Supplementary Figure S2. IGF2BP1 knockdown promotes AJ assembly.** Immunofluorescence stainings of ES-2 (A-C), indicated EOC-derived (D), or none-EOC-derived (E) cell lines transiently transfected by IGF2BP1-directed (siI1) or control (siC) siRNA pools with indicated antibodies, Phalloidin-TRITC staining the F-actin cytoskeleton and DAPI to label the nuclei. Dashed boxes label enlarged regions shown in right panels. Bars 25µM.

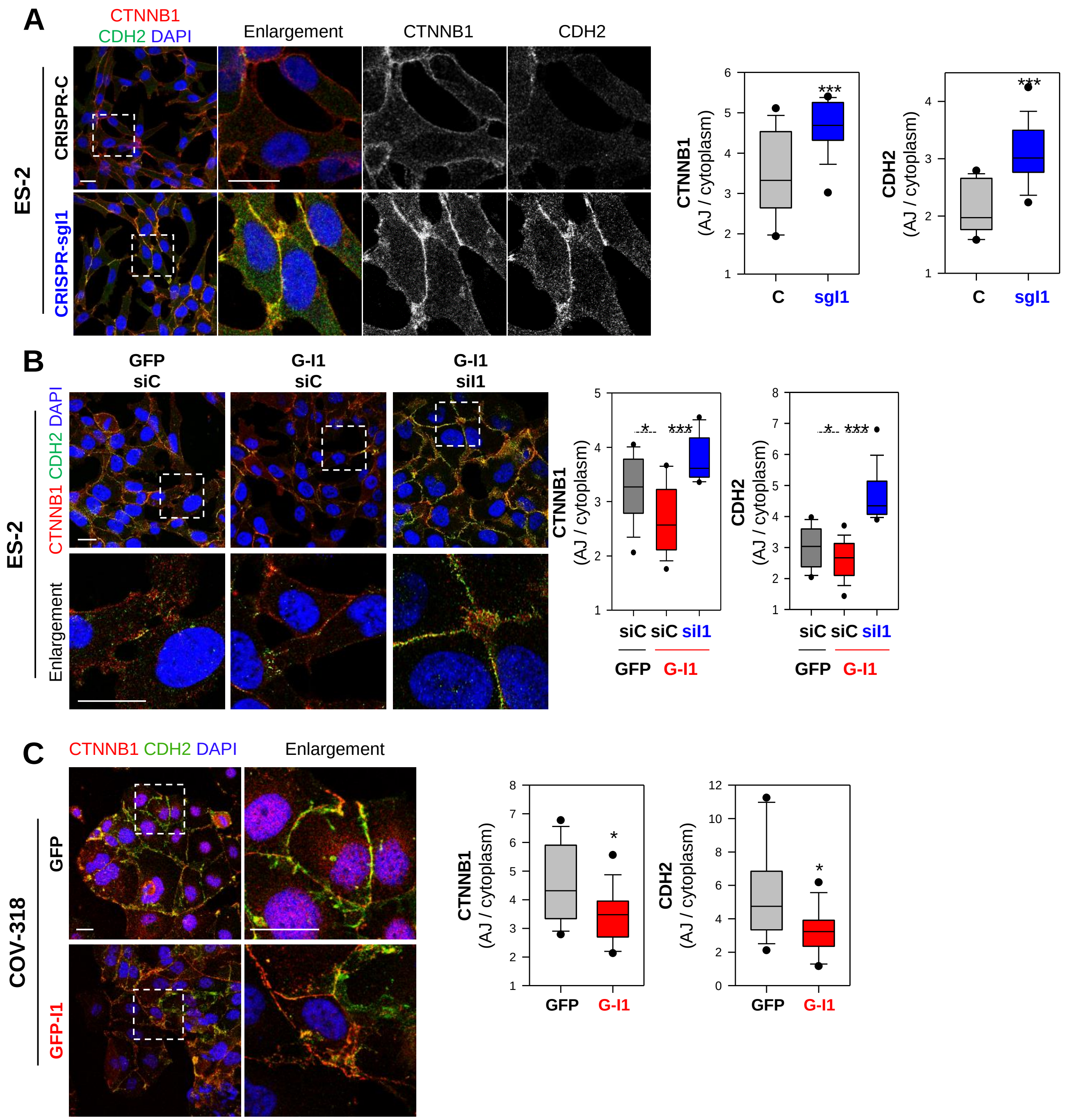


**Supplementary Figure S3. IGF2BP1 promotes AJ disassembly. (A)** Immunofluorescence stainings of ES-2 with CRISPR/Cas9-mediated IGF2BP1 deletion (CRISPR-sgI1, blue) or control (CRISPR-C) cells with indicated antibodies labelling AJ components. Dashed boxes label enlarged regions shown in right panels. Localization of AJ components was quantified at the membrane (AJ) normalized to cytoplasmic staining (box plot, right panel). **(B)** Rescue experiment of ES-2 cells expressing GFP or IGF2BP1. Cells were transiently transfected with indicated siRNAs and analyzed as in (A). IGF2BP1 knockdown restored AJs diminished by IGF2BP1 over-expression. **(C)** COV-318 stably over-expressing IGF2BP1 or GFP were analyzed as in (A). IGF2BP1 over-expression partially releases CDH2 and CTNNB1 from the sites of cell-cell contacts. Bars 25µM.

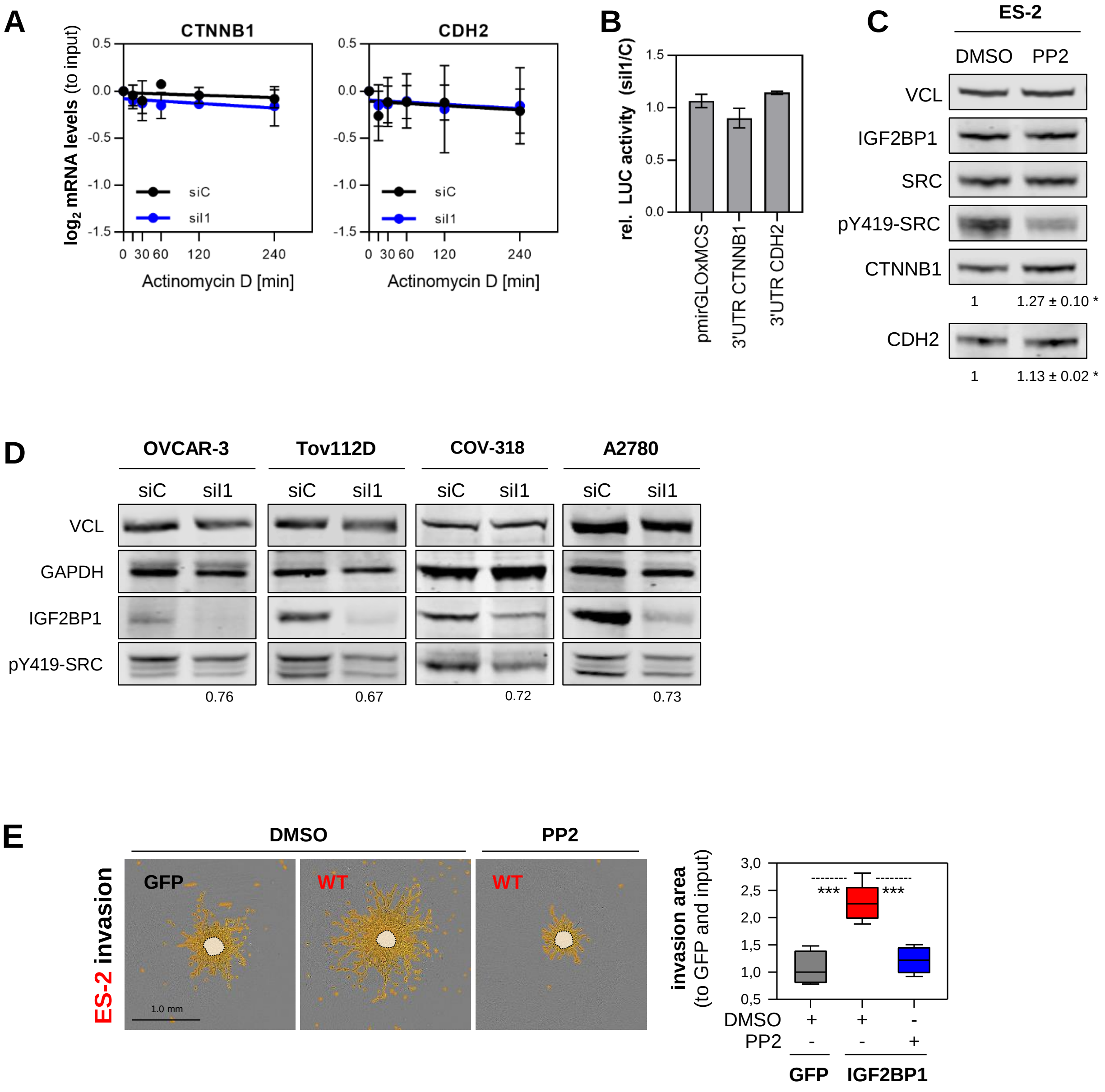
A
CTNNB1
CDH2
log2 mRNA levels (to input)
siC
siI1
Actinomycin D [min]
0 30 60 120 240
0.5 0.0 -0.5 -1.0 -1.5
B
rel. LUC activity (siI1/C)
pmirGLOxMCS
3'UTR CTNNB1
3'UTR CDH2
C
ES-2
DMSO PP2
VCL
IGF2BP1
SRC
pY419-SRC
CTNNB1
1 1.27 ± 0.10 *
CDH2
1 1.13 ± 0.02 *
D
OVCAR-3
Tov112D
COV-318
A2780
siC siI1
VCL
GAPDH
IGF2BP1
pY419-SRC
0.76
0.67
0.72
0.73
E
ES-2 invasion
DMSO
PP2
GFP
WT
WT
1.0 mm
invasion area (to GFP and input)
3,0 2,5 2,0 1,5 1,0 0,5
***
***
DMSO + + -
PP2 - - +
GFP
IGF2BP1

**Supplementary Figure S4. IGF2BP1 controls SRC activity**. **(A)** mRNA turnover of indicated transcripts was monitored by qRT-PCR in ES-2 cells with transient IGF2BP1 depletion (blue) 72h post transfection by blocking transcription using Actinomycin D (5µM) for indicated time points. Quantifications shown $\log_2$ mRNA levels normalized to input (zero time point). RPLP0 served as normalization control. **(B)** Luciferase reporter assays using the 3'UTRs (3' untranslated region) of CTNNB1 and CDH2 fused to firefly luciferase were performed in ES-2 cells with transient IGF2BP1 depletion. A reporter comprising the MCS (multiple cloning site) only served as negative control. Renilla (encoded by the same plasmid) normalized firefly activities relative to the control transfection (siC) are shown. No significant difference between CTNNB1, CDH2 and control reporters was determined. **(C)** Western blot analyses of ES-2 cells treated with PP2 (1µM) or DMSO for 48h using indicated antibodies. Quantifications depicted as number under the respective lanes were determined relative to VCL (C) or total SRC (D) serving as loading control. **(D)** Western blot analyses of indicated EOC-derived cell lines with IGF2BP1 depletion (72h post transfection) using indicated antibodies. Quantifications depicted as number under the respective lanes were determined relative to VCL. **(E)** Spheroid invasion of ES-2 cells with GFP or IGF2BP1 over-expression was monitored using an incucyte S3 device. Representative images overlaid with segmentation masks (yellow) for the invasion or input spheroid (light yellow) are shown. Pre-formed, embedded spheroids were treated with DMSO or PP2 (10µM) as indicated. Qantification of 6 spheroids from three independent experiments is shown as box plot (right panel). Errors represent s.d. from three independent experiments. Statistical significance was determined by Student's T-test. *, $p < 0.05$.

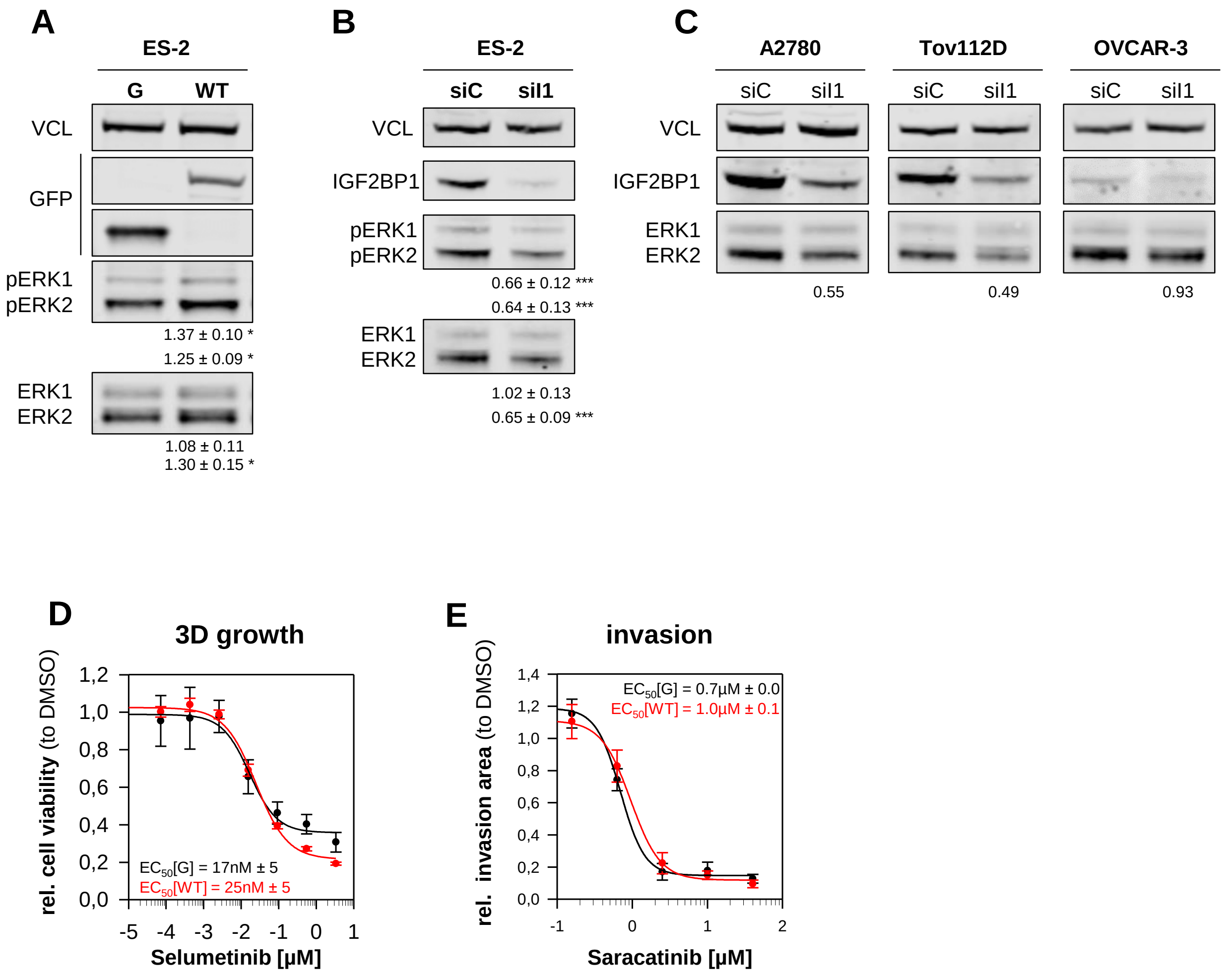


**Supplementary Figure S5. IGF2BP1 controls ERK2 expression. (A-C)** Western blot analyses of IGF2BP1 over-expression (A; G-I1) or depletion (B, C; siI1) with indicated antibodies. VCL served as loading and normalization control for quantifications relative to the control transfections (GFP in A or siC in B) depicted under the respective panels. **(D,E)** EC50 for saracatinib for 3D growth (D) and selumetinib for invasion (E) were determined for indicated inhibitor concentrations (72h). Cell viability was determined using Cell Titer Glo (Promega). Errors represent s.d. (A,B) or s.e. (D,E) from three independent experiments. Statistical significance was determined by Student's T-test. *, $p < 0.05$; ***, $p < 0.001$.

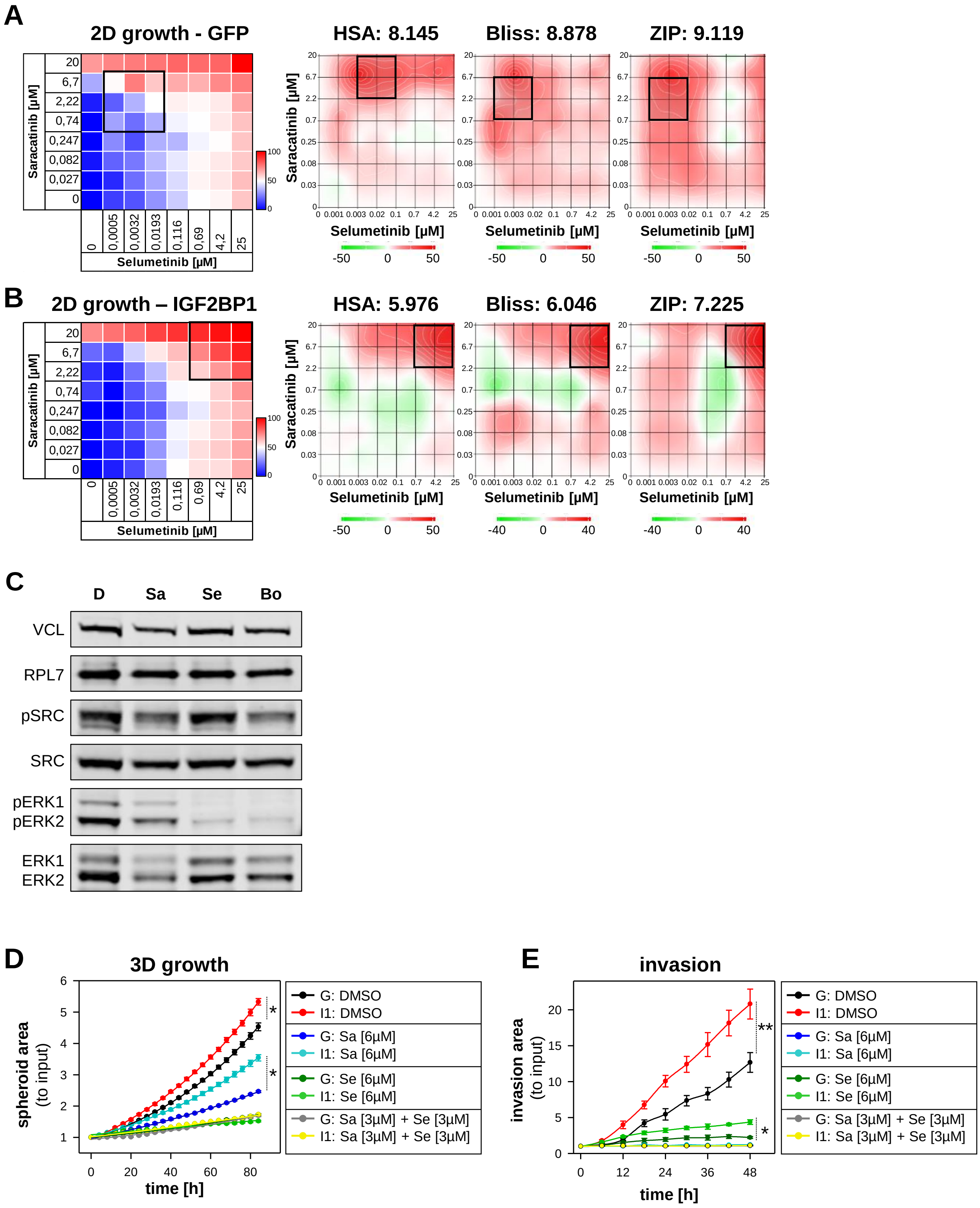
A
2D growth - GFP
HSA: 8.145
Bliss: 8.878
ZIP: 9.119
Saracatinib [µM]
Selumetinib [µM]
B
2D growth – IGF2BP1
HSA: 5.976
Bliss: 6.046
ZIP: 7.225
C
D
Sa
Se
Bo
VCL
RPL7
pSRC
SRC
pERK1
pERK2
ERK1
ERK2
D
3D growth
spheroid area
(to input)
time [h]
E
invasion
invasion area
(to input)
time [h]
G: DMSO
I1: DMSO
G: Sa [6µM]
I1: Sa [6µM]
G: Se [6µM]
I1: Se [6µM]
G: Sa [3µM] + Se [3µM]
I1: Sa [3µM] + Se [3µM]

**Supplementary Figure S6. Additive effects of saracatinib and selumetinib. (A,B)** Synergism between saracatinib and selumetinib was tested using a drug matrix screen (left panel) for GFP or IGF2BP1 over-expressing ES-2 cells under 2D growth conditions with indicated concentrations of the respective drugs for 72h. Cell viability was determined by Cell Titer Glo (Promega). Inhibition is shown in (%) as heat map (left panel). Synergy maps were accomplished using https://synergyfinder.fimm.fi/ (5), identifying additivity for both conditions (synergy score > -10; < 10) using indicated models. Synergy scores are shown as color-coded contour plots (scores < -10, antagonistic, green; scores > 10, synergistic, red). HSA, highest Single Agent model; Bliss, Bliss Independence model; ZIP, Zero Interaction Potency model. Notably, concentrations for maximal additivity concentrations (indicated as black frame) are higher for IGF2BP1 compared to GFP, confirming resistance. **(C)** Western blotting with indicated antibodies was used to confirm effective SRC and ERK inhibition. (Sa: 6µM saracatinib; Se: 6µM selumetinib; Bo: 3µM saracatinib and 3µM selumetinib). **(D,E)** Time resolved graphs to Figure (6E-H). Notably, spheroid growth is diminished but not abolished by saracatinib and invasion, although reduced, still occurs upon selumetinib treatment. Error depicts s.e. from three different experiments. Statistical significance was tested by Student's T-test. *, $p < 0.05$; **, $p < 0.01$.

### Supplementary Figure Legends

**Supplementary Table T1A. IGF2BP1 mRNA expression in TCGA-OV-RNA-Seq cohort.** Table shows the TCGA patient identifyers, the $\log_2$ (RSEM) expression data for IGF2BP1 and its classification in high ($\log_2$ (RSEM) > 5) or low ($\log_2$ (RSEM) < 5) based on the TCGA-OV-RNA-Seq dataset.

**Supplementary Table T1B. Local HG-SOC tumor cohort.** Table summarizes our local ovarian cancer cohort consisting of fresh frozen and FFPE samples. IGF2BP1 mRNA expression data determined by RNA sequencing as well as the Remmele score determined by immunohistochemistry staining (IHC) using IGF2BP1-directed antibodies are indicated.

**Supplementary Table T2. Differential gene expression of IGF2BP1 high vs low expressing tumors.** Table summarizes the official gene symbols and differential gene expression between samples with high and low IGF2BP1 expression (Supplementary Table T1) for each gene.

**Supplementary Table T3. GSEA for positive enriched gene sets.** Gene set enrichment analyses was used to determine positively enriched HALLMARK gene sets in samples with high and low IGF2BP1 expression. Differential gene expression (Supplementary Table T2) was used for gene ranking.

**Supplementary Table T4. Classification of EOC cell lines.** RNA-Seq data obtained from the CCLE project were used to classify EOC cell lines as epithelial-like or mesenchymal-like by their mRNA expression of epithelial (CDH1, EPCAM, KRT8) or mesenchymal (CDH2, VIM, ZEB1) markers. Median expression was used as cut-off. Four cell lines (nd) did not fall into either category. Models used in this work are labeled in bolt.

**Supplementary Table T5. Quantitative proteomics.** Extracts of IGF2BP1 depleted or control cells (5 replicates) were TMT labeled and analyzed in an Orbitrap Fusion Tribrid mass spectrometer (Thermo Fisher). Quantifications based on the reporter ion ratios derived from high-resolution MS/MS spectra is shown for each replicate.

**Supplementary Table T6. Cell lines.** All cell lines and corresponding RRID identifiers, distributors and catalogue numbers are summarized as table.

**Supplementary Table T7. Antibodies.** Antibodies, companies, catalogue numbers and RRID identifiers are shown for all primary and secondary antibodies used in this study.

**Supplementary Table T8.** Plasmids and oligonucleotides. Table summarizes all plasmids generated in this study including oligonucleotides and restrictions sited used for cloning. Primer pairs used for qRT-PCR are listed as well.

### Supplementary Video Legends

**Supplementary Video V1: IGF2BP1 promoted 3D growth is abolished by combined SRCi/MEKi treatment.** Pre-formed spheroids of GFP (control; uppel panel) or IGF2BP1 over-expressing ES-2 cells (lower panel) were treated with indicated inhibitors: saracatinib [6μM]; selumetinib [6μM] or combination [3μM saracatinib + 3μM selumetinib]. 3D growth was monitored over time in an Incucyte S3 (Sartorius). Growth area was determined by spheroid segmentation shown by red overlay. Videos are related to Figure 6.

**Supplementary Video V2: IGF2BP1 promoted invasive potential is abolished by combined SRCi/MEKi treatment.** Pre-formed spheroids of GFP (control; uppel panel) or IGF2BP1 over-expressing ES-2 cells (lower panel) were imbedded into Matrigel ® and treated with indicated inhibitors: saracatinib [6μM]; selumetinib [6μM] or combination [3μM saracatinib + 3μM selumetinib]. Invasion was monitored over time in an Incucyte S3 (Sartorius). Invasive growth (indicated by overlayed segmentation mask in red) was quantified. Videos are related to Figure 6.
